# NucleosomeDB - a database of 3D nucleosome structures and their complexes with comparative analysis toolkit

**DOI:** 10.1101/2023.04.17.537230

**Authors:** Grigoriy A. Armeev, Anna K. Gribkova, Alexey K. Shaytan

**Affiliations:** Department of Biology, Lomonosov Moscow State University, 119234 Moscow, Russia; Department of Computer Science, HSE University, 109028 Moscow, Russia

**Keywords:** nucleosome structure, chromatin structure, chromatin proteins, PDB

## Abstract

Nucleosomes are basic building blocks of chromatin, comprising DNA wrapped around an octamer of eight histone proteins. They play key roles in DNA compaction, epigenetic mark up of the genome and actively participate in chromatin dynamics. X-ray and later cryo-EM studies have contributed greatly to our understanding of nucleosome structure, their interactions and dynamics. With over 470 nucleosome containing structures in the Protein Data Bank there is a wealth of information to be gleaned from these structures, especially through comparative analysis. However, due to the variability in their representation (chain naming, residue numbering, other artifacts), these structures cannot be systematically analyzed “as is”. To address this issue, we developed a framework for analyzing and classifying nucleosome structures and their complexes, resulting in the creation of the NucleosomeDB database and the corresponding web-service. NucleosomeDB allows researchers to search, explore, and compare nucleosomes with each other, despite differences in composition and peculiarities of their representation. By utilizing the information contained within the NucleosomeDB, researchers can gain valuable insights into how nucleosomes interact with DNA and other proteins, assess the implications of mutations and protein binding on nucleosome structure. The detailed information contained within NucleosomeDB can contribute to a better understanding of the structure and function of nucleosomes, and ultimately, the functioning of chromatin and gene regulation. NucleosmeDB is freely available at https://nucldb.intbio.org.

## Introduction

Genomic DNA in eukaryotes, together with a set of nuclear proteins, is stored in chromatin. Chromatin provides packaging of genetic material, allowing for transcription, replication, and DNA repair processes to occur. At the level of chromatin, numerous mechanisms of epigenetic regulation of gene transcription are implemented, which significantly affect gene expression and, as a result, control the functioning of cells and organisms. The organization of chromatin determines the degree of DNA accessibility for transcription factors, polymerases, histone chaperones, and many other proteins that directly control the processes of gene expression, cell differentiation, and influence the processes of cellular aging.

At the very first level of organization, the packaging of eukaryotic DNA is carried out by forming nucleosomes - particles with a size of about 10 nm. The compact part of the nucleosome - the nucleosome core particle (NCP) consists of approximately 145-147 base pairs of DNA wrapped around a hetero-octamer of histones of four types [1]. Groups of nucleosomes form chromatin fibers and ultimately allow DNA to be packed into dense chromosomes that form during cell division.

However, the specific structure of intermediate levels of organization is unclear. Currently, the concept of chromatin organization in interphase cells has moved away from the classical model, in which nucleosome fibers form 30-nanometer fibers with subsequent packaging of such fibers into higher order fibers [2]. Moreover, it has been shown that chromatin is largely similar to polymer solutions and demonstrates microphase separation [3,4]. The regulation of chromatin function is carried out by a huge number of regulatory proteins, a significant part of which can bind to histones or DNA within the nucleosome. Moreover, nucleosomes themselves demonstrate significant variability in composition and properties, they can consist of various non-canonical histone variants [5]. Additionally, histones can undergo post-translational modifications (PTMs) that affect transcription levels [6].

Currently, the Protein Data Bank (PDB [7]) includes more than 470 structures containing nucleosomes. These structures shed light on the details of interactions with regulatory proteins, changes caused by the introduction of PTMs, and mechanisms of chromatin compaction at the oligonucleosome level [5,8–10]. However, all these structures were accumulated over the past 25 years, resulting in significant variability in chain naming and residue numbering. Structures contain artifacts, and cannot be subjected to systematic analysis as is. By analyzing the structures of nucleosomes, we can gain insight into how they interact with DNA, how they are modified, and how they are positioned within chromatin. Understanding the structures of nucleosomes is a critical step in understanding how they regulate gene expression and other cellular processes. For this reason, we have developed a framework for analyzing and classifying nucleosome structures and their complexes, and implemented it in the form of NucleosomeDB. Our database allows searching, exploring, and comparing nucleosomes with each other, despite differences in composition, naming conventions and residue numbers.

## Materials and methods

### Data extraction and preprocessing

The current database update was performed on 08.03.2023 and is based on PDB data available at the respective date. We extracted data from RCSB PDB using RCSB PDB Data API in several steps.

1. We used the curated set of histone sequences from HistoneDB [11] to query the matching polymer instances in the PDB via the RCSB PDB REST API. We found 711 unique PDB IDs by queuing 303 different histone sequences. Several structures out of 711 matched the same polymer entities to the different histone types and thus were excluded, leaving 693 non-redundant histone-containing structures.
2. Based on the structure metadata, we filtered the dataset to collect structures that contain at least one pair of H3-H4 histones and at least one double-stranded DNA segment, which yielded 481 structures for further analysis. All these structures were downloaded for structural preprocessing and annotation. For all structures, we analyzed biounits. Two structures were not analyzed due to their size as they could not be converted to the standard PDB format (8GXQ,8GXS), several structures were converted to PDB format (7UNC, 7UND, 7XSE, 7XSX, 7XT7, 7XTD, 7XSZ, 7XTI,7YWX). All analysis was performed with author provided chain names and residue numbers, as automatic renumbering and renaming provided by RCSB is inconsistent.
3. All structures were analyzed using custom scripts written on python with MDAnalysis [12] library. On this step, we detected the DNA type and histone variant with BLAST using the curated set of DNA sequences (see Supplementary table 1). We assigned DNA strands as top and bottom in the way they were defined in the curated set, in case of palindromic sequences chains were defined in alphabetical order. We used pairwise sequence alignment to match histone sequences to canonical sequences from the structure with PDB ID 1KX5, which allowed us to map histone fold components, such as α-helices.
4. We detected all nucleosome-like particles and subnucleosomes using several geometrical thresholds.
  a. Histone dimers were detected using 10 Åthreshold between centers of masses (COM) of histone fold Cα atoms.
  b. H3H4-H3H4 tetramers were detected using 16.8 Åthreshold between COMs of H3 α3 α-helices (thus forming a 4-helix bundle).
  c. H3H4-H2AH3B tetrameres were detected using 20 Åthreshold between COMs of H4 and H2B α3 α-helices.
5. We then classified nucleosome-like particles into different categories: octasomes, hexasomes and hemisomes. Afterward, we calculated the number of said particles in the structures (biounits). In this analysis, we did not check if different histone cores are located on the same DNA strand. Therefore, those structures contain not only multinucleosome structures but also separate nucleosomes, which were assigned to the same biounit.

On this step the database contained 479 structures with the meta information about the structure, extracted IDs of proteins.

### Structure analysis

To simplify further analysis and visual inspection, we placed all structures in the common nucleosome reference frame (NRF). The NRF is determined by the superhelical axis of nucleosomal DNA (Z-axis), the pseudo symmetry dyad axis (Y-axis), and the vector product of Y-axis and Z-axis (X-axis). This procedure was performed using structural alignment of histone folds with the reference structure, which was put in the NRF by the procedure described in [13]. Histone folds were mapped by pairwise sequence alignment to the canonical histones using BioPython [14].

We performed the analysis of DNA geometry for all structures. To simplify further analysis, we detected the nucleotide base pair for which C1′ atoms are closest to the dyad axis (see Figure 2 B). Then we assigned basepairs starting from the dyad using geometrical and sequence criteria. We only analyzed the structure of double-stranded DNA belonging to the first nucleosome in the structure. For all basepapairs we then analyzed DNA geometry using 3DNA software and calculated the relative twist of nucleosomal DNA as described in [13].

#### Contacts analysis

To analyze and compare the molecular interactions within nucleosomes and their complexes with other proteins, we performed contact analysis. We searched for atom pairs located closer than 4 Åto each other and belonging to different residues and polymer chains, and summed the number of such interactions to create interaction profiles for every polymer chain in the structure.

#### DNA geometry analysis

We assigned pairs of nucleotides based on complementarity of the upper and lower DNA strands sequences. This approach proved more stable than structural methods for determining pairs in this case, as the space of available DNA sequences in nucleosomes is limited. To compare analysis results, we numbered each pair of nucleotides relative to the pair closest to the nucleosome’s dyad axis. We chose the numbering direction to match the direction of numbering of the upper DNA strand. We analyzed DNA geometry for nucleosomal DNA using 3DNA [15] and we also calculated the “relative twist” for each DNA basepair [16]. This parameter varies from −180 to 180 degrees, but we plot the absolute value of the parameter for ease of visualization. All DNA parameters were renumbered dyadwise.

We determined the extent of DNA unwrapping based on the analysis of DNA geometry of aligned structures. We considered a DNA segment (starting from either end of the nucleosomal DNA) to be unwrapped from the NCP if the position of every base pair in that segment (as provided by 3DNA) was more than 7 Åaway from the position of any base pair in the reference 1KX5.

#### Histones geometry comparison

We analyzed all octasomes that contained all four types of histones. All structures were pre-aligned to the reference structure using the histone folds. To compare the structures with each other, we mapped all residues in the structures to the reference structure using pairwise alignment of histone sequences (these alignments are available in “compare alignments” tool). We only selected those amino acids that could be mapped in all structures, thus creating atomic selections that contained amino acids that had equivalents in all other structures (common atom selections). In further analysis, we only used the coordinates of Cα atoms. We calculated the radii of gyration and standard deviations of the atom coordinate components for each structure based on common atom selections. For ease of visualization, we also performed PCA on coordinate deviation components.

We compared all structures with each other using the interatomic distance matrices calculated from common atom selections. We calculated the root-mean-square deviations of interatomic distances for all pairs of structures. This approach avoids artifacts associated with aligning all structures to the reference structure. Using the described metric, we clustered all structures using the hierarchical agglomerative clustering method (as implemented in Scipy [17]) with a distance cutoff of 1 Å.To visualize the clusters, based on the difference matrix between structures, we constructed a dendrogram (http://nucldb.intbio.org/struct_phylo_tree).

### Interacting proteins’ classification

To classify nucleosome interactors, we designed empirical classification of chromatin proteins (based on Gene Ontology [18]) and then classified non-histone proteins according to PDB extracted protein name, UniProt functional annotation and manual curation. Classification consists of 4 major classes - Function (Chromatin remodeling, DNA modification, Histone PTM erasers, Histone PTM readers, Histone PTM writers, Histone chaperones, Kinetochore, RNA polymerases, Synthesis of second messengers, Transcription factors), Process (Antiviral defense, DNA repair, DNA replication, Nuclear division, Transcription-associated), Genomic location (Centromere-associated, Telomere-associated), Viral (DNA integration, Viral).

### Database construction

As NucleosomeDB combines structural data with protein and DNA sequences with metadata, we chose not to use the classical database approach and instead used a set of JSON objects with a common dictionary. We chose such an approach to simplify the development, as by its nature, NucleosomeDB is a moderated and limited subset of specific structures from the PDB. Data stored in serialized objects is then accessed through the web application written in Flask (https://github.com/pallets/flask). For the visualization, we use NGL JavaScript library [19], and for plotting - combination of D3 via the python interface MPLD3 with custom plugins to visualize sequence features and alignments. The database is deployed on a dedicated server of the research group.

## Results

We created a framework for the extraction, annotation and analysis of the structures of nucleosomes available in the PDB. Using this framework we constructed a web database NucleosomeDB which provides tools for data exploration and analysis. NucleosomeDB contains 479 structures of nucleosomes, complexes of nucleosomes and supranucleosomal structures. We annotated, aligned, analyzed all structures and prepared collections of structures on the landing page. On Figure 1A we superimposed all structures, containing a single nucleosome, to demonstrate the overall structure variety. Since 2017, an increasing number of nucleosome structures have been obtained using the cryo-EM method (Figure 1B). In 2022, all structures (a record number, over 80) were obtained using this method. The application of cryo-EM significantly expanded the set of known complex structures (Figure 1C), but at the same time, many structures with relatively low resolution appeared (Figure 1B).

**Figure 1.**
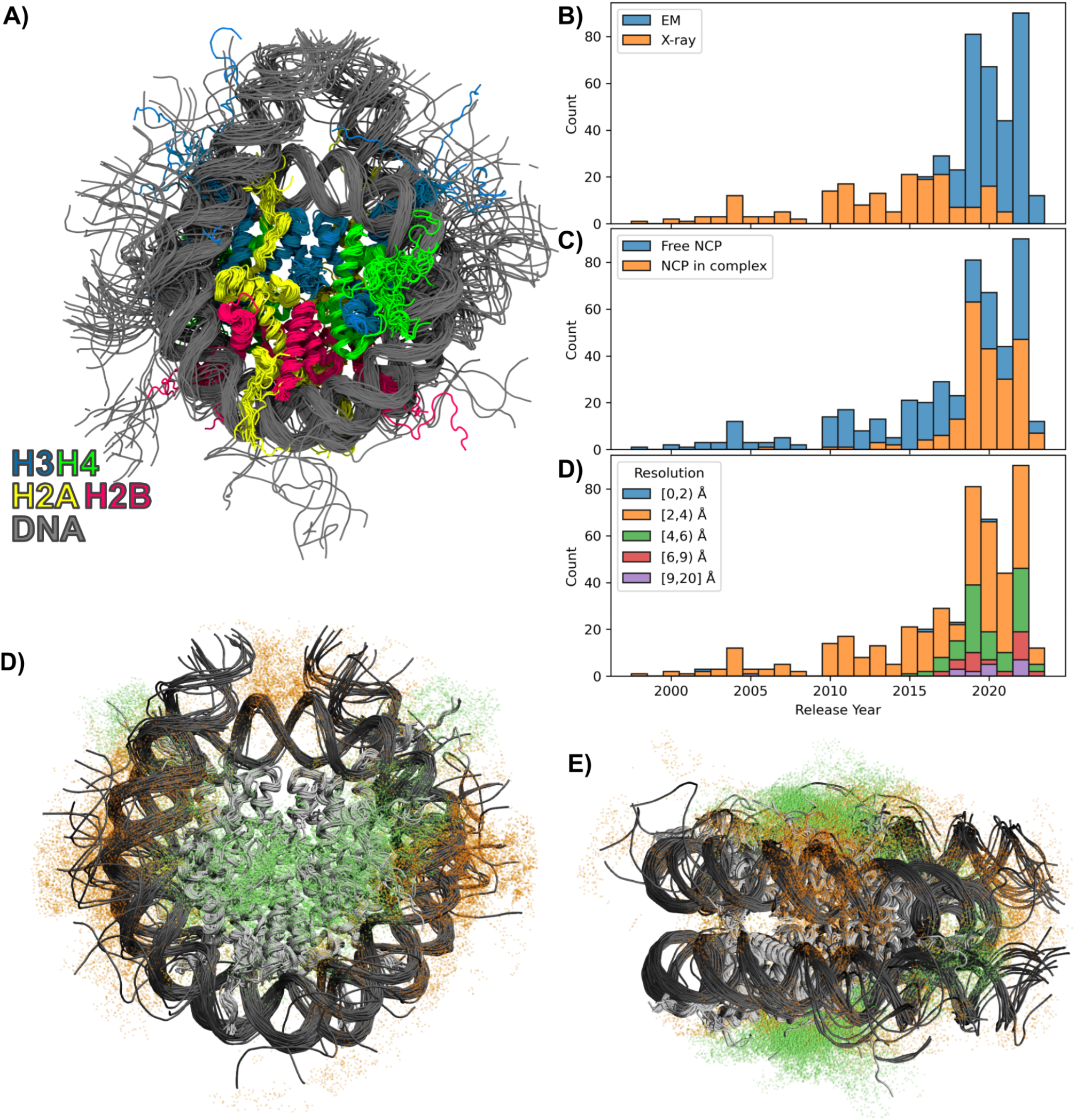
A) Overlay of all structures of individual nucleosomes in NucleosomeDB. Histones and DNA are shown as tubes, interacting proteins are hidden. B)-D) Distribution of structures in the database by year of release depending on the method, structure type, and resolution. D) E) Ensemble of atoms from proteins interacting with nucleosomes. Green dots depict proteins interacting with histones, orange dots depict proteins interacting with DNA.

**Figure 2.**
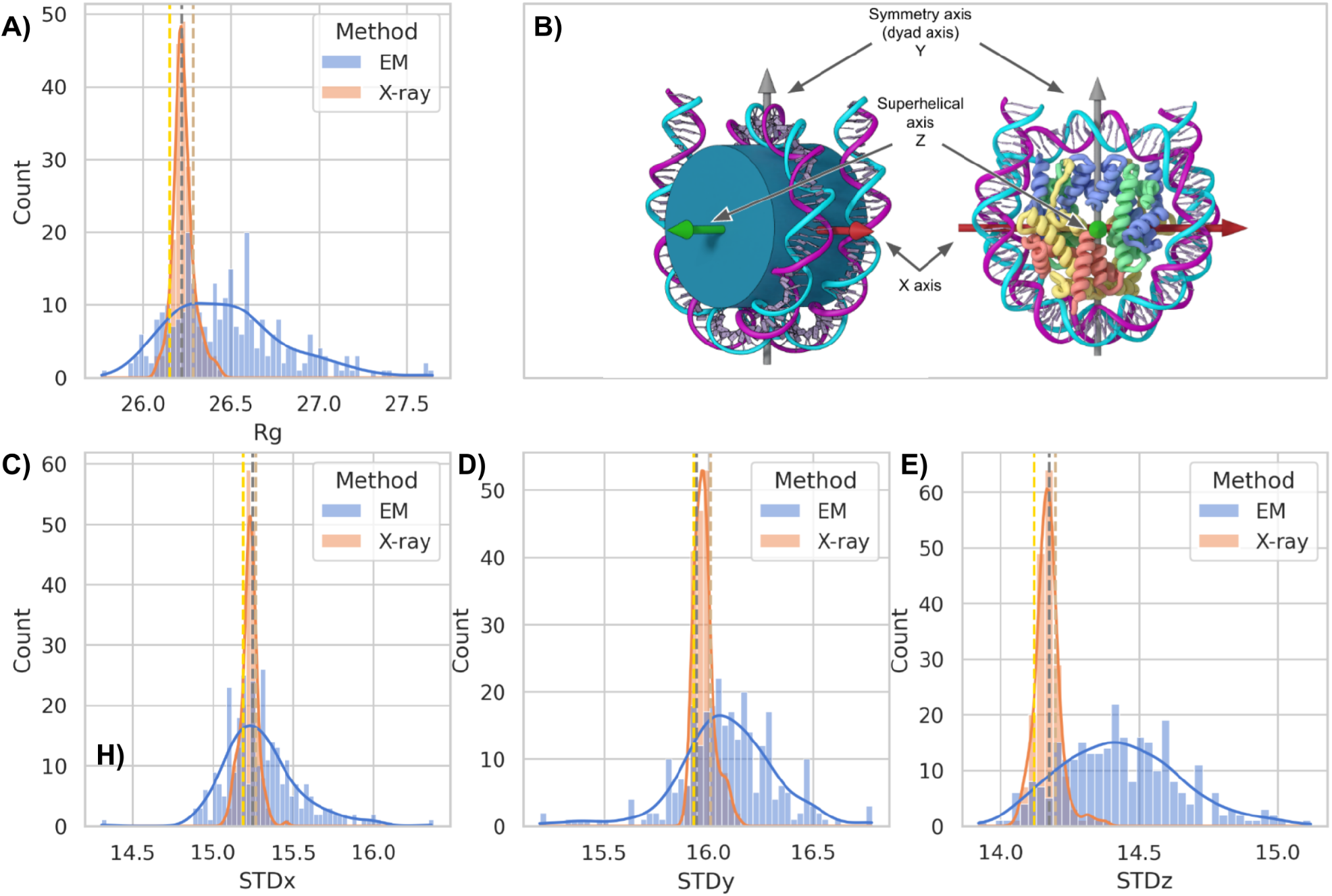
Figure 1. A) Distribution of radius of gyration for the overall set of atoms for all nucleosome structures (see methods) depending on the method of structure determination. B) Visualization of nucleosome reference frame (NRF) and nucleosome axes direction. C)-D) Distribution of standard deviation of atomic coordinates by NRF components.

NucleosomeDB is a web resource, which allows users to explore the variability of histone variants and DNA within the nucleosomes, browse different nucleosome-containing complexes and supranucleosomal structures. Below is a brief overview of the contents of the database by categories: histone variants, DNA positioning sequence variants, variability of nucleosome interactors, variability of supranucleosomal structures

Histone variants play crucial roles in gene expression, DNA replication, and DNA damage repair. These variants have unique amino acid sequences, which affect nucleosomes ability to interact with DNA and other proteins. Even a few amino acid substitutions in histone variants can alter nucleosome dynamics and stability, leading to changes in gene expression and other cellular processes. The H3 histone type is the most represented in nucleosome structures, the PDB contains nucleosomes with H3.3, cenH3, TS H3.4, H3.5, H3.6, and H3.Y. Similarly, H2A variants such as H2A.1, H2A.B, H2A.J, H2A.X, H2A.Z, are also present in nucleosome structures. MacroH2A is also present but only in truncated form. There is only one H2B.1 variant present in nucleosome structures. Nucleosomes in database contain human and model organisms histones: *Homo sapiens* (219 structures), *Xenopus laevis* (172), *Saccharomyces cerevisiae* (12), *Drosophila melanogaster* (8), *Mus musculus* (7). But there are structures of nucleosomes with histones of *Komagataella pastoris* [20], nucleosomes of pathogenic unicellular eukaryotic parasite *Giardia lamblia* [21] and human nucleosome with incorporation of protozoan parasite Leishmania histone H3 [22]. Archaeal nucleosomes are not included in the database due to incompatibility with data processing pipeline. The linker histone H1 binds to the nucleosome, forming what is known as a chromatosome. There are a total of 33 structures containing linker histones, with the most predominant variant being human H1.4 (20 structures). Additionally, NCP structures contain human H1.0 and H1.10 variants, chicken H.5, and one structure from *Xenopus* containing H1.0 and H1.8.

While there is no direct readout of DNA sequences during nucleosome formation, it is known that the stability and positioning of nucleosomes depend on the nucleosomal DNA sequence. The nucleosomes in PDB can be classified into three main groups based on their DNA sequences. The first group contains the most structures and includes the Widom 601 sequence [23] and its derivatives - 246 structures. The second sequence group includes α-satellite sequences and their derivatives - 166 structures (natural sequence from human centromeric region). The third sequence group includes 10 structures with telomeric sequence. There are also several other DNA sequences including cen3 DNA, MMTV, DNA1, sat2 and Human-D02 and unclassified. Despite the fact that PDB contains hundreds of different nucleosome structures, their DNA sequences belong to a limited pool of sequences and their variations.

There are 193 structures containing NCPs in complex with other proteins. The majority of NCP complexes with non-histone interactors belong to the following biological processes: 4 major classes - Function (10 categories), Process (5), Genomic location (2), Viral (2). We also provide a list of curated keywords to find all related structures. We focused this work on the analysis of structures diversity, thus we did not analyze the nucleosome interactom, however we provide tools for comparative contacts analysis. The largest number of structures are associated with transcription and DNA repair (Process class) and relate to histone PTM writers, chromatin remodelers and transcription factors (Function class). We also classified structures by genomic location, with 21 and 7 structures associated with centromere and telomere regions, respectively. A more detailed analysis of nucleosome complexes with non-histone protein diversity and its cancer relation has been published at [24]. The repertoire of available structures is quite broad, we superimposed all structures with interacting proteins with resolution better than 4 Å and showed all atoms from the interacting proteins within 10 Å of histones and DNA. From the (Figure 1 DE), it can be seen that the entire surface of the histone octamer is uniformly surrounded by interacting protein atoms.

However, it is quite difficult to identify preferred binding regions, and this requires further study. Interestingly, there are regions of DNA that are completely free of known interacting protein structures (Figure 1 DE).

We also analyzed all supranucleosome structures (structures containing two or more nucleosomes). There are 24 such models, both with and without a linker histone. Here, we assessed the available models (bioassemblies) and counted nucleosomes located on continuous double-stranded DNA. The majority of supranucleosomal structures contain two nucleosomes (11 structures), 8 structures contain three nucleosomes and 5 structures contain four nucleosomes and one structure contain six nucleosomes (PDB ID: 6HKT).

### Structure variability

We investigated the distribution of the radius of gyration of the protein component of nucleosomes for all structures containing a complete set of histones (see methods). Interestingly, the radius of gyration for the most compact structure was 25.76 Å, while for the largest it was 27.65 Å.The most frequently occurring radius of gyration was 26.22 Å (Figure 2 A). For the reference structure (1KX5), the radius of gyration was 25.15 Å. Thus, protein regions of nucleosomes differ in size from the reference in the range from -1.5% to 5.7%. It is particularly interesting that variations in sizes are weakly dependent on structure resolution, but significantly dependent on the method of structure determination. We observed a narrow distribution of radius of gyration for structures determined by XRD. On the other hand, structures determined by the EM method showed a significantly broader distribution with a shift of the distribution center (Figure 2 A, 26.48 for EM and 26.2 for XRD).

The radius of gyration is a way to measure how much molecules are different in size (given that we compare equivalent atoms). Such a metric is not suitable for determining unevenly deformed structures. Since in our case, all molecules were pre-aligned, we compared the distribution of deviations of atomic coordinates (Figure 2 CDE). The distributions of coordinates deviations along X and Y were unimodal and fluctuated within a fairly narrow range. However, the distribution of atomic deviations along the Z axis was significantly broadened. As with the radius of gyration, structures obtained by cryoEM demonstrated a broadened and shifted distribution. Average cryoEM structures are deformed in Y and Z axes (Figure 2 DE). To simplify the selection of deformed structures, we conducted principal component analysis for distributions of atomic coordinates. The first principal component describes the uniform change in the size of the molecule (Figure 3 B), and the two remaining components describe deformation of the nucleosome (Figure 3 C). The figure shows that most structures are deformed uniformly, but there are N structures that are unevenly deformed (https://nucldb.intbio.org/struct_variety).

**Figure 3.**
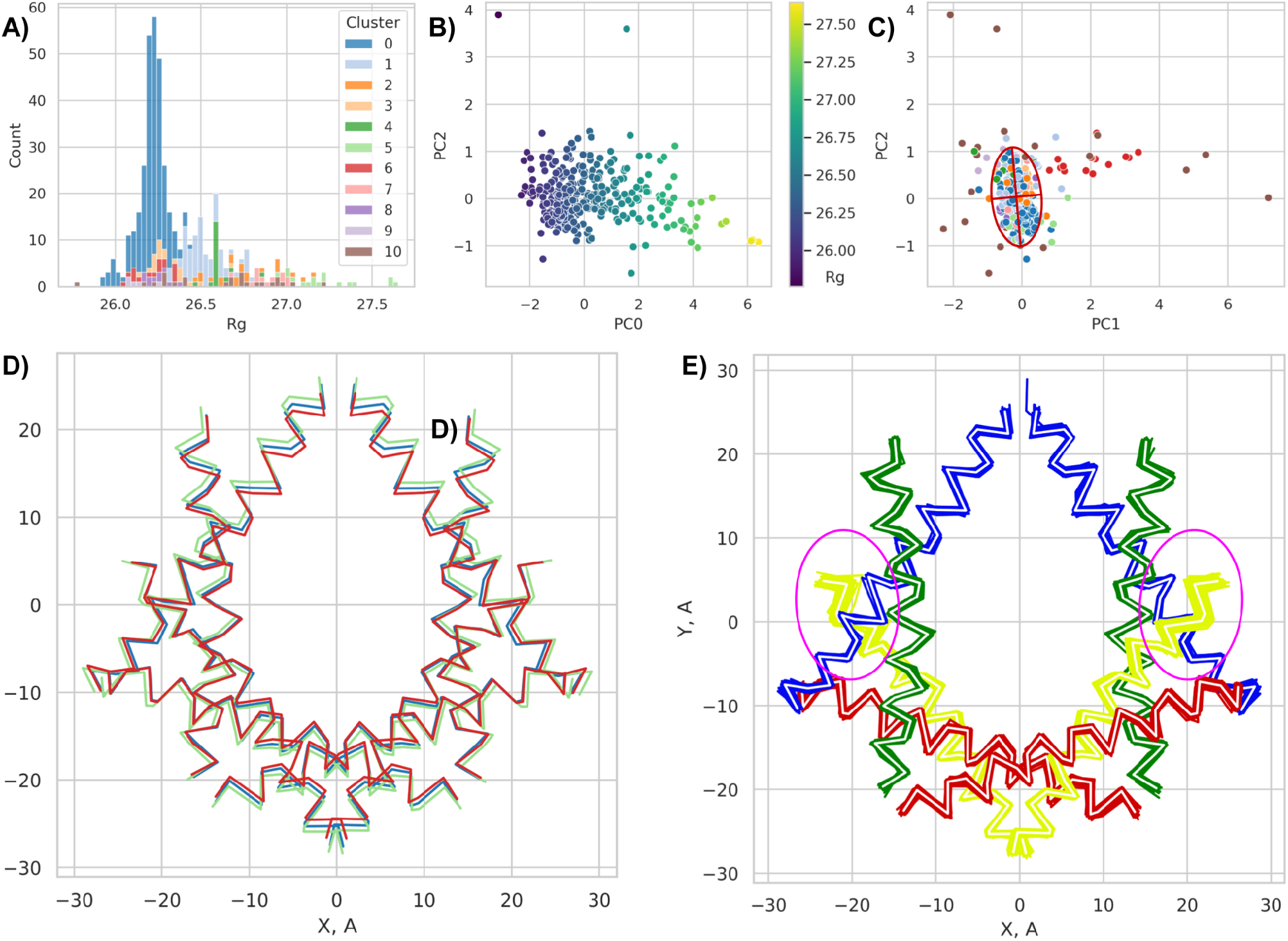
A) Distribution of radii of gyration colored by the structure cluster. B) and C) Projections of standard deviation of atomic coordinates on principal components. D) Projections of averaged α2 helices of the four clusters (color scheme from A). E) Projections of α2 helices from all structures within the first structure cluster.

We performed a cluster analysis for all nucleosome structures based on pairwise comparison of their interatomic distance matrices. As a result, we identified 18 clusters of structures, with half of them consisting of two or fewer structures (Supplementary table 2, https://nucldb.intbio.org/struct_phylo_tree). Thus, we merged all small clusters into cluster number 10. More than half of the known structures fell into one structural cluster, indicating high similarity within these structures (root-mean-square deviation of interatomic distances for structures within the cluster is less than 1 Å). This cluster predominantly includes compact structures obtained by the XRD method, with an average resolution in this cluster of (Figure 3 A). The second most populated cluster consists mainly of structures obtained by the EM method. The remaining clusters consist almost exclusively of EM structures. The (Figure 3 D) shows the difference in the averaged geometry for the 4 most populated clusters. Those clusters mainly differ in Y direction - the direction of symmetry axis. We also observed an increased variety in histone H2B helices positions inside the most populated class with very similar structures (Figure 3 E).

We evaluated the distribution of structures with partially unwound DNA from histones. We assessed the degree of unwinding in two ways. 1. For structures containing enough DNA to form complete nucleosomes, we evaluated the differences in the positions of the nucleotide pair centers compared to the reference structure. We found 41 structures in the PDB with this type of unwinding (Figure 4 A). Projections of DNA traces for such structures are shown on Figure 4 BD. Such DNA unwinding is often associated with nucleosome binding to partner proteins and does not describe the possible mobility of DNA itself (only 9 of 41 are free nucleosome structures). The dynamics of DNA unwinding are difficult to study using structural methods; in X-ray crystallography, all mobility is limited by crystalline packing, and in cryo-EM maps, there are no areas with increased mobility because they are averaged from multiple images. To assess the possible average degree of DNA unwinding using cryo-EM, we selected structures with unmodeled DNA segments, although the structure annotation indicates that full-size DNA was used to prepare the sample (69 structures 11 from which free are free nucleosomes). The resulting length distribution indicates the possibility of unwinding DNA up to 30 base pairs from the nucleosome entry point and up to 50 base pairs from the nucleosome exit point (Figure 4 C). Such degrees of unwinding are also associated with the formation of nucleosome complexes with other proteins.

**Figure 4.**
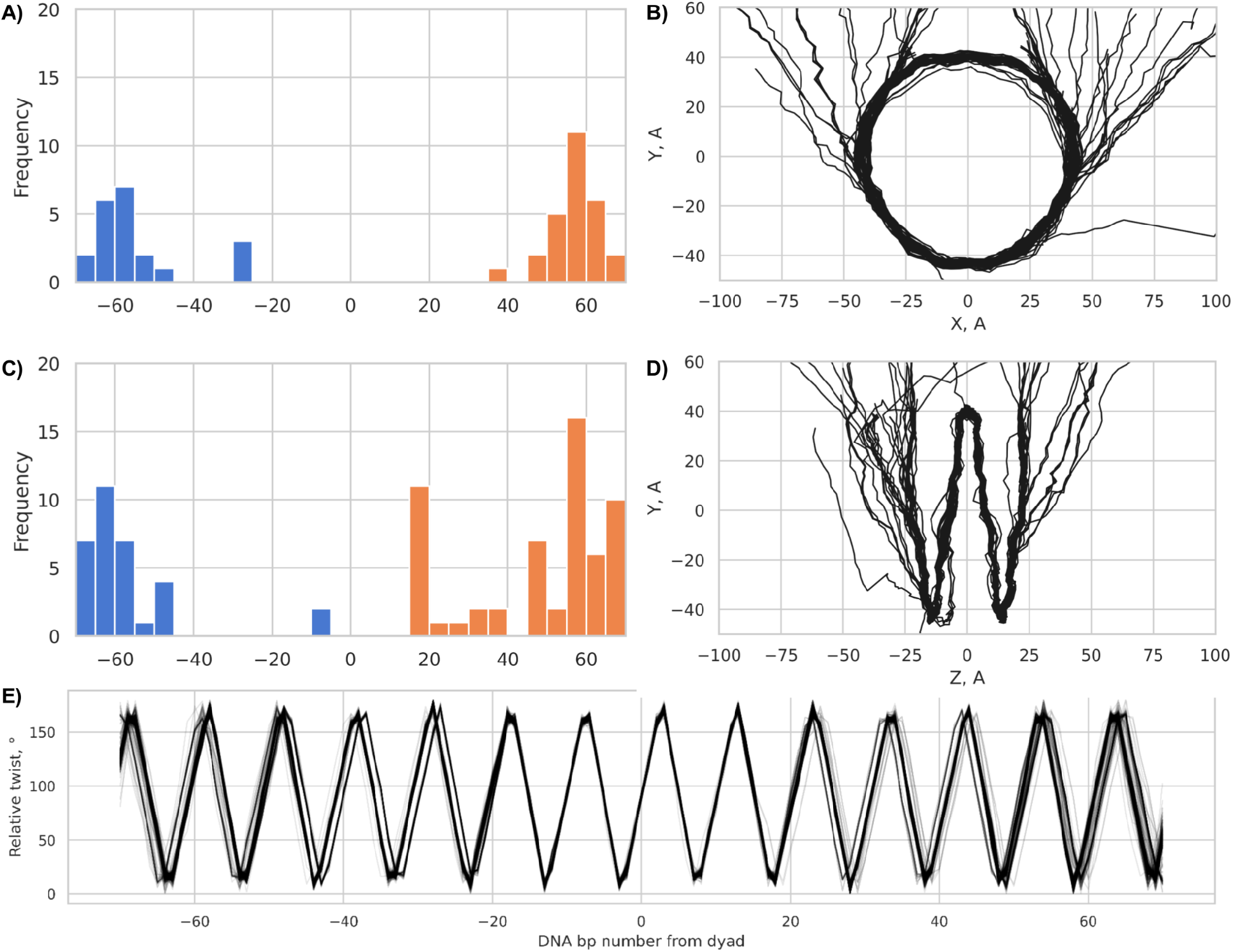
A) Distribution of structures with partially unwrapped DNA. B) D) Projections of all structures with partially unwrapped DNA. C) Distribution of cryoEM structures with unmodeled DNA E) DNA Relative twists for all structures that contain at least 141 b.p. of DNA

We also calculated the relative twist values for all nucleosomes that contain at least 70 nucleotides from both sides of the closest to dyad base pair (370 structures meet this requirement). For this analysis, we only used structures without register shifts. As can be seen from the (Figure 4 E), the density of supercoiled DNA wrapping can vary for DNA with equivalent lengths. In the regions of entry and exit of the nucleosome, three possible orientations of the edge nucleotide pairs are distinguishable. Interestingly, there is almost no variability in relative twist in the region from -20 to +20 from dyad base pair.

### Database web interface

NucleosomeDB is publicly available at https://nucldb.intbio.org/ without registration. The user can browse structures, see available Statistics, Search structures and download individual aligned structures. All the metadata and structural features data is available via the API described in Help section. This section also contains usage examples and methods description. The web interface also allows users to add several structures to the collection for the later analysis and comparison.

Each structure has its own page that displays the composition of the structure with annotations of histone variants, DNA, and nucleosome-interacting proteins. The page includes a visualization of the structure with the option to compare it with the reference. Annotation tables also allow highlighting individual chains in the visualization. All pages feature a button to download the structure placed in a nucleosome reference frame, as well as three tabs with interactive graphs: a structure projection graph for assessing deformation visually, a DNA geometry tab containing Relative Twist graphs and 3DNA parameters, and graphs of nucleosome component contacts.

Contact graphs allow visualizing interactions in the structure window.

On individual structure pages, as well as on the Browse page and on the search results page, buttons are available to add structures to the user’s collection. The collection tool enables users to compare projections of multiple structures to explore differences in DNA and histone geometry between structures, and compare DNA-contact profiles with proteins. Additionally, users have access to a page for three-dimensional visualization of superimposed structures, along with alignment of histone component sequences. This interface enables determination of the impact of changes in histone and DNA sequences.

## Discussion

We developed a framework for analyzing and classifying nucleosome structures and their complexes, which we implemented in the form of NucleosomeDB database and web-service to aid in the analysis of nucleosome structures. Our database allows searching, exploring, and comparing nucleosomes with each other, despite differences in composition and nomenclature. To our knowledge, this is the first specialized database that provides access to a curated and preprocessed set of nucleosome strucures. The structures of nucleosomes provide valuable insights into how they interact with DNA, how they are modified, and how they are positioned within chromatin.

NucleosomeDB provides a framework for the systematic analysis of nucleosome structures and their complexes.

We found that the radius of gyration of the protein component of nucleosomes varies from -1.5% to 5.7% compared to a reference structure (1KX5). The size variations are not dependent on structure resolution, but significantly dependent on the method of structure determination (Figure 2, https://nucldb.intbio.org/struct_variety). The sizes of nucleosome structures can vary more in cryoEM since there are not constrainted by the crystalline environment. Several studies have shown that the protein core of the nucleosome is not flexible, with the plasticity of the core being linked to the conformational mobility of DNA and the binding of other proteins that interact with nucleosomes. However, such a significant scatter of sizes may also indicate possible imperfections in cryoEM data processing methods. In cryoEM, the equivalent pixel size may be determined imprecisely, as its determination requires regular calibration. To improve it, the exact voxel size can be selected by performing a solid-state fitting of homologous structures to the experimental electron density map. Flexible fitting of a new structure is carried out only after such scaling. However, this step is not a standard or mandatory part of the data processing pipeline. As a result, molecular models may be uniformly enlarged or shrunken. A uniform increase in the model by several percents does not lead to molecular clashes or other noticeable defects in the structures. We showed that the quality of the models improves from year to year for the set of nucleosomes while resolution is not improving [24]. This indicates possible overfitting of the models to the experimental maps.

We found that structures obtained by cryoEM showed a broadened and shifted distribution of atomic deviations along the Z and Y axis, indicating some structures are deformed unevenly (Figure 2 CDE). The possibility of nucleosome deformation has been discussed for quite some time, different modes of nucleosome mobility have been distinguished, one of which affects the histone core of the nucleosome [25]. In particular, the broadening of the nucleosome structure along the Z axis (the axis of the nucleosome superhelix) can be attributed to gaping motions. The role of deformed nucleosomes is also discussed in the work [26], in which such deformation is associated with the process of DNA unwinding. Interestingly, the relative variability of the geometry of nucleosomes along the Z axis is higher than along the X axis. This phenomenon can be explained by molecular constraints imposed by the tightly wound DNA loop around the histone octamer.

CryoEM structures are also deformed along the Y axis compared to the XRD structures. Such non-uniformity may be a consequence of the data analysis. It is known that in cryoEM, nucleosomes tend to point in the same direction when frozen in ice. In most images, the nucleosomes are aligned parallel to the detector plane. Such non-uniformity leads to anisotropic blurring in Z direction and other angle dependent artifacts inside electron density maps. The y-axis is a C2 symmetry axis in nucleosome, thus some artifacts may be introduced by particle alignment algorithms. However, for automatic annotation of such artifacts, a stable robust approach to data processing and deposition of intermediate processing results is necessary.

We conducted a cluster analysis of all nucleosome structures and found that 9 classes of structures can be distinguished. The root-mean-square deviation (RMSD) between any two structures within the class was no more than 1 Å (https://nucldb.intbio.org/struct_phylo_tree). Additionally, the database contains 24 structures that are significantly deformed and do not fall into these clusters. Interestingly, the affiliation to a cluster depends on the method of structure determination, as the first cluster predominantly consists of data obtained by NMR or cryoEM structures derived from NMR structures. The eighth cluster consists entirely of structures obtained by NMR and containing variant histones H3 and H2A. Thus, it should be noted that despite the large number of structures, the variability within them is quite low. However, even within clusters with similar structures, internal plasticity can be observed. When superimposing all structures from the first cluster, we found that the ends of the H2A histone alpha helices have a greater coordinate spread (Figure 3 E) than the other histones, which is consistent with molecular modeling data [27].

We discovered that DNA models in cryoEM structures can have unmodeled base pairs from both sides (up to 30 base pairs from the nucleosome entry point and up to 50 base pairs from the nucleosome exit point, Figure 4 AC). DNA unwrapping from nucleosomes was studied with a variety of methods [16,28–30]. Although it cannot be directly detected in electron density maps, electron density smearing suggests high mobility and unwrapping. It is interesting that we observed some asymmetry in DNA unwrapping profiles, but as we did not consider differences in DNA sequences, this observation may be misleading. Also, due to nucleosome symmetry, there may be some artifacts introduced by cryoEM data analysis during particle alignment.

### Limitations and updating plans

At present, our database does not contain structures of archaeal nucleosomes or nucleosomes with viral histones due to significant differences in histone sequence. The database also does not contain structures of individual histone dimers and their complexes. Adding all of these structures is planned in future updates of the database. In the analysis of structures, alternative locations of atoms were not used. For structures containing more than one model, only the first model was analyzed. For structures with multiple nucleosomes, only the first one appearing in the structural file was analyzed. Structures 8GXQ and 8GXS contain nucleosomes but were not included in the database as the number of chains in them currently exceeds the capabilities of the structure processing pipeline.

In the future, we plan to expand the database and analyze interactions between nucleosomes and other proteins by mapping interactions of individual histones. We also plan to introduce metrics for evaluating model quality based on local resolution maps for cryo-EM structures.

## Conclusion

The study of nucleosome structures is crucial in understanding how genetic information is stored, accessed, and regulated. To facilitate this study, various databases have been developed that provide information about nucleosome positioning and composition and possible interacting proteins. We created a database that provides currently available structural information in a unified manner. We have demonstrated that the given set of known interactors fully covers the globular regions of histones and DNA areas in the SHL2 region. Furthermore, we have shown that nucleosomes significantly differ from each other in size, not always uniformly, which may indicate both an internal plasticity and possible artifacts in the processing of experimental data. We have also discovered a wide range of DNA unwinding variants and different modes of DNA supercoiling, which requires further investigation.

In conclusion, NucleosomeDB can aid in the identification and characterization of nucleosome-associated proteins and the study of nucleosome dynamics and interactions with other molecules. We hope that this tool will allow researchers to gain a deeper understanding of the mechanisms of gene regulation and develop new therapies for diseases that involve alterations in chromatin structure and function.

## Acknowledgements

We thank Andrey Moiseenko for valuable comments about the cryoEM data analysis pipelines.

The research was carried out using the equipment of the shared research facilities of HPC computing resources at Lomonosov Moscow State University.

