## Supplementary information for "NucleosomeDB - a database of 3D nucleosome structures and their complexes with comparative analysis toolkit"

**Supplementary table 1**. DNA sequences used for classification

| Name | Length | Dyad position(s) | doi |
|---|---|---|---|
| **601** | 232 | 133 | 10.1080/073911010010524947 |
| **601L** | 146 | 73 | 10.1093/nar/gks261 |
| **601_modified** | 146 | 73 | 10.1126/science.aau9904 |
| **MMTV** | 438 | 114,311 | 10.1080/073911010010524947 |
| **alphasat_147** | 147 | 73 | 10.1016/S0022-2836(02)00386-8 |
| **alphasat_146** | 146 | 73 | 10.1016/S0022-2836(02)00386-8 |
| **alphasat_146_4CpG** | 146 | 73 | 10.1002/2211-5463.12064 |
| **alphasat_146b** | 147 | 73 | 10.1016/S0022-2836(02)00386-8 |
| **alphasat_native** | 145 | 72 | 10.1038/s41467-019-10247-4 |
| **alphasat_A16** | 146 | 73 | 10.1016/j.jmb.2004.07.080 |
| **alphasat_4_palindorm** | 147 | 73 | 10.1038/nature10258 |
| **alphasat_2L** | 145 | 73 | 10.1098/rsob.150128 |
| **alphasat_2R** | 146 | 73 | 10.1098/rsob.150128 |
| **DNA-1** | 148 | 74 | 10.1038/s41586-020-2195-y |
| **Human-D02** | 146 | 73 | 10.1038/s41467-019-12007-w |
| **telomeric_human** | 146 | 73 | 10.1101/2019.12.18.881755 |

***NucleosomeDB - a database of 3D nucleosome structures and their complexes***

***supplementary information***

Supplementary table 2. Structure PDB IDs with respective average structure parameters.

| Cluster | PDBID | Rg | STDx | STDy | STDz |
|---|---|---|---|---|---|
| 0 | 7K5Y, 8H1T, 7K63, 6LA2, 6LA8, 6LA9, 6LAB, 7COW, 7K5X, 5NL0, 4QLC, 5WCU, 6L9Z, 7K60, 1EQZ, 3AV1, 4KGC, 4LD9, 5B0Z, 6DZT, 8GPN, 2CV5, 3AFA, 3AZG, 3AZI, 3AZJ, 3AZK, 3AZL, 3AZM, 3AZN, 3W96, 3W97, 3W99, 3WKJ, 4YM5, 4YM6, 5AV5, 5AV6, 5AV8, 5AV9, 5AVB, 5AVC, 5B1M, 5B24, 5B2I, 5B2J, 5CPI, 5CPJ, 5CPK, 5GSE, 5GTC, 5JRG, 5XF3, 5XF4, 5XF5, 5Y0C, 5Y0D, 6HKT, 6JR0, 6JR1, 6K1I, 6K1J, 6K1K, 6KVD, 6L49, 6L4A, 6LER, 6M3V, 6M44, 6T93, 6USJ, 6V2K, 6V92, 6YOV, 7C0M, 7D1Z, 7K61, 7LYC, 7NL0, 7SCY, 7SCZ, 2NQB, 2PYO, 3AYW, 3AZE, 3AZF, 3AZH, 6FML, 6VZ4, 8AV6, 3AV2, 3LEL, 3WTP, 3X1S, 3X1T, 3X1U, 3X1V, 5B0Y, 5E5A, 5GSU, 5GT0, 5GT3, 5X7X, 5XM0, 6J99, 6LTJ, 6NE3, 6RYR, 7UNC, 7UND, 5B40, 1KX3, 1KX4, 1KX5, 1S32, 1ZBB, 2FJ7, 2NZD, 3B6F, 3B6G, 3C1B, 3KUY, 3LJA, 3LZ0, 3LZ1, 3MGP, 3MGQ, 3MGR, 3MGS, 3MNN, 3MVD, 3O62, 3REH, 3REI, 3REJ, 3REK, 3REL, 3TU4, 3UT9, 3UTA, 3UTB, 4J8U, 4J8V, 4J8W, 4J8X, 4R8P, 4WU8, 4WU9, 4XUJ, 5B1L, 5CP6, 5DNM, 5DNN, 5OXV, 5OY7, 5XF6, 5XM1, 6NZO, 6O96, 6OM3, 6PA7, 6PX1, 6WZ5, 6WZ9, 6X0N, 7E8D, 7E8I, 7OH9, 7OHA, 7OHB, 7OHC, 7XPX, 8G86, 8G88, 8G8B, 8G8G, 3A6N, 3KXB, 6ZHX, 7JZV, 7U51, 7U52, 1ZLA, 6VYP, 3C1C, 5F99, 5GXQ, 5OMX, 7PGV, 7PGW, 7PH5, 7PH6, 7ZS9, 7ZSA, 7ZSB, 8CEO, 1M18, 1M19, 1M1A, 1P3B, 1P3F, 1P3G, 1P3I, 1P3O, 1P3P, 5ONG, 5ONW, 1P34, 1P3A, 1P3K, 1P3M, 1P3L, 4KUD, 4Z5T, 4JJN, 1ID3, 7DBH, 5Z23, 5ZBX, 5AY8, 1AOI, 3W98, 6IPU, 6IQ4, 6JM9, 6JMA, 6JXD, 7CCQ, 7CCR, 4XZQ, 4YS3, 4Z66, 6Y5D, 7K6P, 7K6Q, 3KWQ, 5MLU, 6Y5E, 6KE9, 6L9H, 6LE9, 4X23, 3AN2, 6E0C, 6E0P, 6SEG, 7D20, 7PII, 7YYH | 26.21 | 15.2 | 15.97 | 14.17 |
| 1 | 8H0V, 8H0W, 7KBF, 5KGF, 5X0X, 6PWE, 6PWF, 7KBD, 7KBE, 6R8Y, 6R8Z, 6R90, 6R91, 6R92, 6R93, 6R94, 6T90, 7LYA, 7LYB, 7XZX, 7XZZ, 7Y00, 5X0Y, 7EG6, 7ENN, 8ATF, 6RYU, 7VDT, 7VDV, 7VVU, 7VVZ, 7W9V, 7XSE, 7XSX, 7XSZ, 7XT7, 7XTD, 6T7D, 6UGM, 6UH5, 5Z3U, 5Z3V, 6ESF, 6JYL, 6R1U, 6R25, 6TDA, 6UXW, 7A08, 7MBN, 6X59, 6X5A, 7OTQ, 7U50, 7U53, 6NJ9, 6S01, 6VEN, 7SWY, 7TN2, 7VBM, 7BXT, 7Y7I, 7K78, 7YQK, 7XD1, 6R1T, 7KTQ, 7XD0, 6IY2, 6IY3, 7XCR, 7XCT, 7WLR, 6KXV, 6O1D, 7U46, 7U47 | 26.52 | 15.31 | 45276 | 14.45 |
| 2 | 6KW5, 7V96, 6R0C, 7BWD, 6IRO, 6K1P, 6KIV, 6NN6, 6NOG, 7MBM, 6NQA, 6BUZ, 6C0W, 6MUO, 6MUP, 6SE0, 6SE6, 6SEE, 6SEF, 6TEM, 7U4D | 26.76 | 15.46 | 16.26 | 14.58 |
| 3 | 7PET, 7PEU, 7PEZ, 7PFT, 7PFU, 7PFV, 7PFW, 7PFX, 4ZUX, 6PWV, 6PWW, 6PWX, 7Y8R, 7PEV, 7PEW, 7PEY, 7AT8, 7M1X, 7BY0 | 26.31 | 44972 | 15.99 | 14.33 |
| 4 | 5O9G, 6A5L, 6A5O, 6A5P, 6A5R, 6A5T, 6A5U, 6INQ, 6IR9, 6J4W, 6J4X, 6J4Y, 6J4Z, 6J50, 6J51, 5Z3L, 5Z3O, 7K7G, 6QLD | 45103 | 44972 | 16.23 | 14.59 |
| 5 | 7DBP, 7JO9, 7JOA, 7TAN, 6KIU, 6KIW, 6KIX, 6KIZ, 7UD5, 7CRO, 7CRP, 7CRQ, 7CRR, 7UV9, 6UPH, 7R5R, 7YWX | 27.24 | 15.81 | 45062 | 14.83 |
| 6 | 7PEX, 7PF0, 7PF2, 7PF3, 7PF5, 7PF6, 7PFA, 7PFC, 7PFD, 7PFE, 6T79, 6T7A, 6T7B, 6T7C, 7PF4, 7PFF, 6FQ5 | 26.22 | 15.36 | 15.63 | 45030 |
| 7 | 7V90, 7V9C, 7V9J, 7V9K, 7V9S, 7VA4, 6T9L, 6GEJ, 6GEN, 7EA5, 7EA8, 7ON1 | 26.88 | 15.52 | 16.31 | 14.69 |

| | | | | | |
|---|---|---|---|---|---|
| 8 | 2F8N, 1U35, 3WA9, 3WAA, 5B31, 5Z30, 6JOU, 5B32, 5B33, 1F66 | 26.24 | 15.18 | 45001 | 44971 |
| 9 | 6W5I, 6W5M, 6W5N, 7E9C, 7E9F | 26.13 | 14.96 | 15.96 | 14.28 |
| 10 | 6KW3, 6KW4, 6HTS, 6M4D, 6M4G, 6M4H, 6UPL, 7XZY, 6G0L, 7EGP, 6I84, 6RNY, 7NKX, 7NKY, 6ESG, 6ESH, 6ESI, 6XJD, 8DU4, 6FQ6, 6FQ8, 6FTX, 6Z6P, 7D69 | 26.62 | 15.38 | 44942 | 14.58 |
